## Supplementary Figures for "iPHoP: an integrated machine-learning framework to maximize host prediction for metagenome-assembled virus genomes"

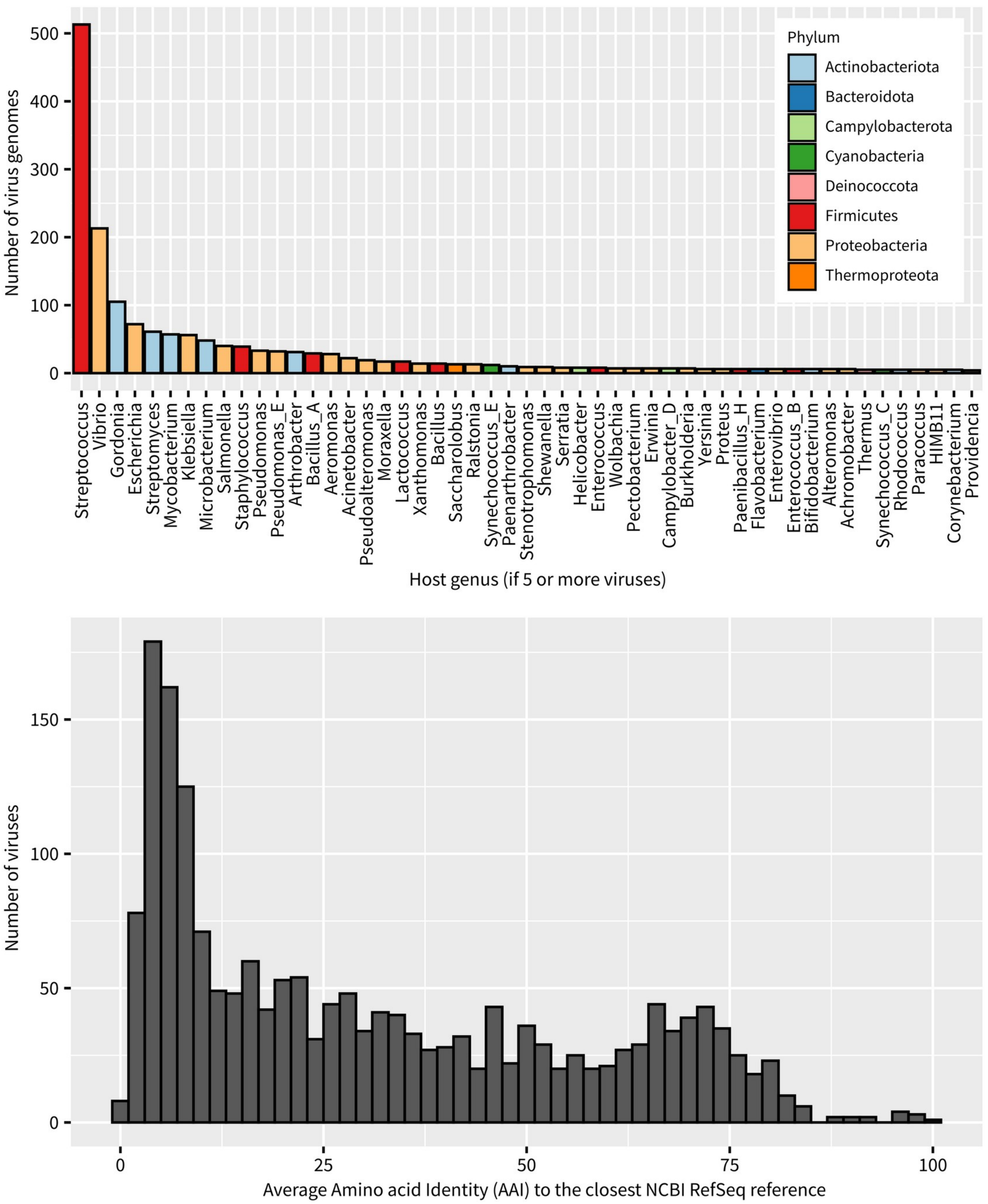


**Supplementary Figure S1. Characteristics of the test dataset.** A. Distribution of the host genera for the test dataset. Note: only genera associated with ≥ 5 viruses are included, another 125 host genera were associated with < 5 viruses and are not displayed. B. Distribution of AAI to the closest reference in NCBI RefSeq for the test dataset. The corresponding list of viral genomes included in the test dataset is provided in Table S2.

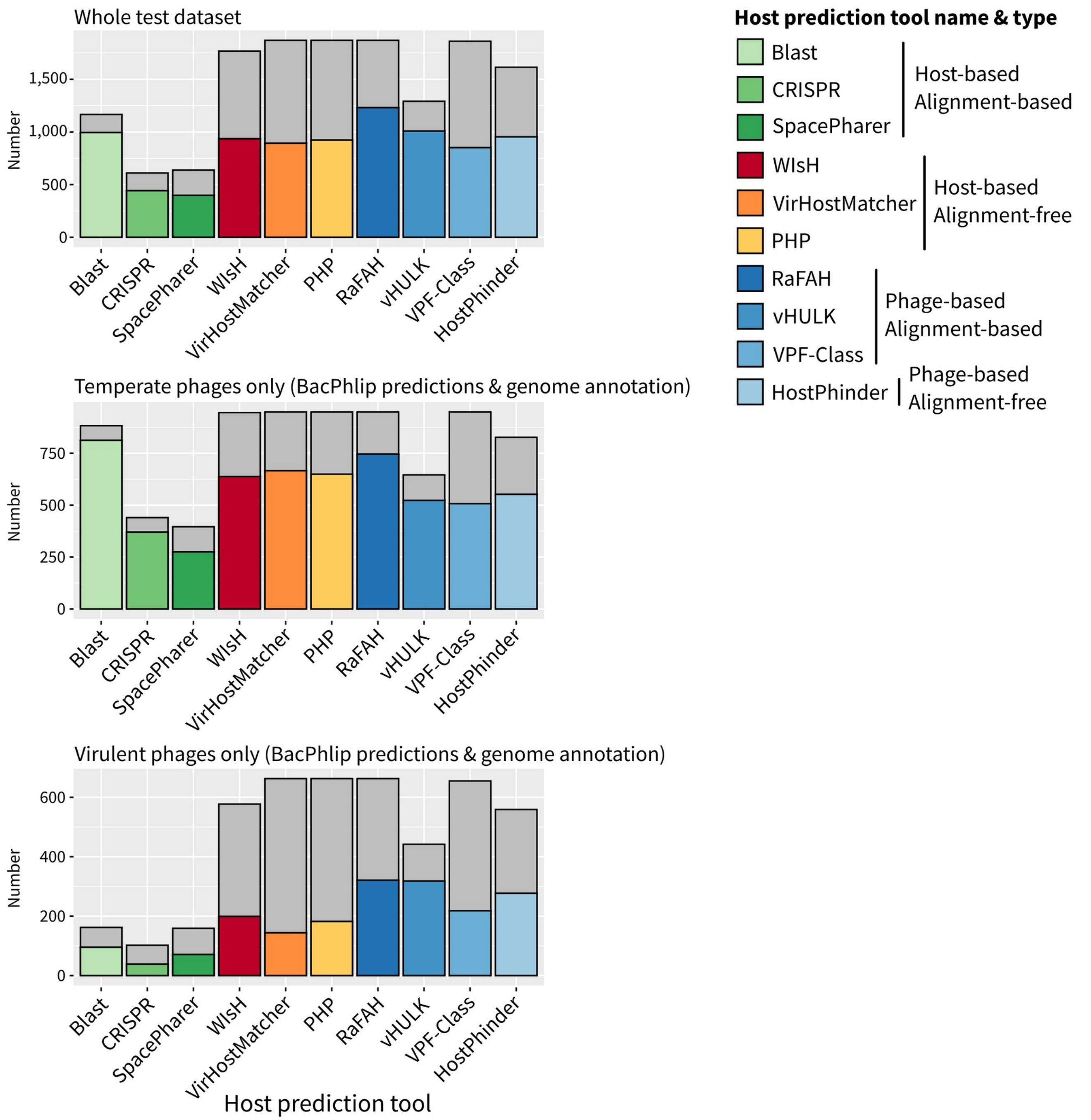


**Supplementary Figure S2. Comparison of different host prediction approaches on different subsets of the test dataset.** Total number of predictions and number of correct predictions (y-axis) obtained at any rank for each tool (x-axis) on sequences from the test dataset (Table S2). For each tool, the number of correct predictions is indicated by the colored bar, while the total number of predictions is indicated by the gray bar. The top panel displays the results obtained on the entire test dataset (n=1,870). The middle panel includes results obtained for all phages predicted as temperate, either via BacPhlip or based on the genome annotation (n=949). The middle panel includes results obtained for all phages predicted as virulent by BacPhlip (n=663).

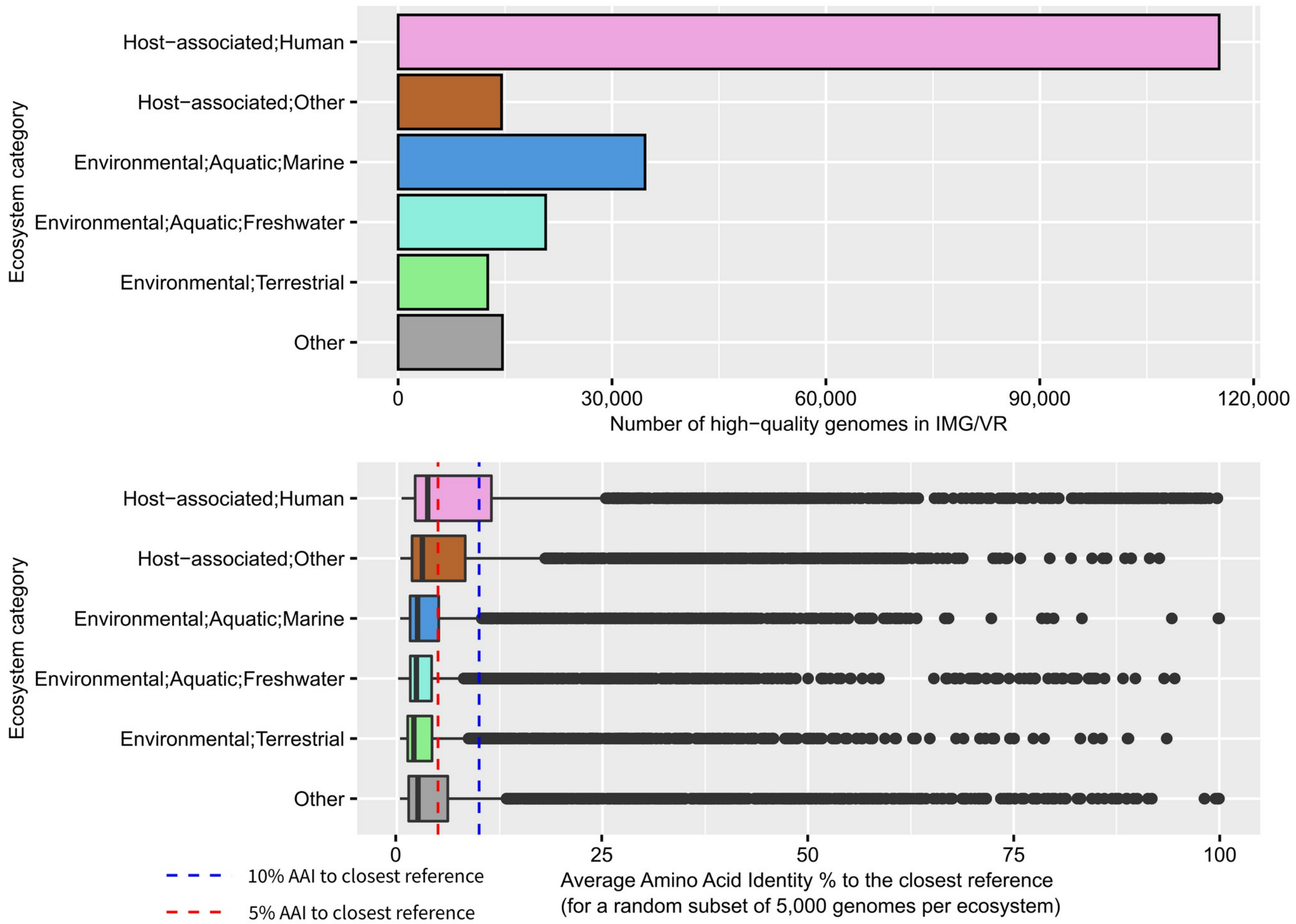


**Supplementary Figure S3. Characteristics of the high-quality IMG/VR genomes**. A. Number of high-quality viral genomes from IMG/VR v3 identified across the 5 major biomes in the database. Genomes sampled from other biomes of lacking a biome information are gathered in the "Other" category. B. Distribution of the average amino-acid identity between IMG/VR v3 viral genomes and the NCBI Viral RefSeq v203.

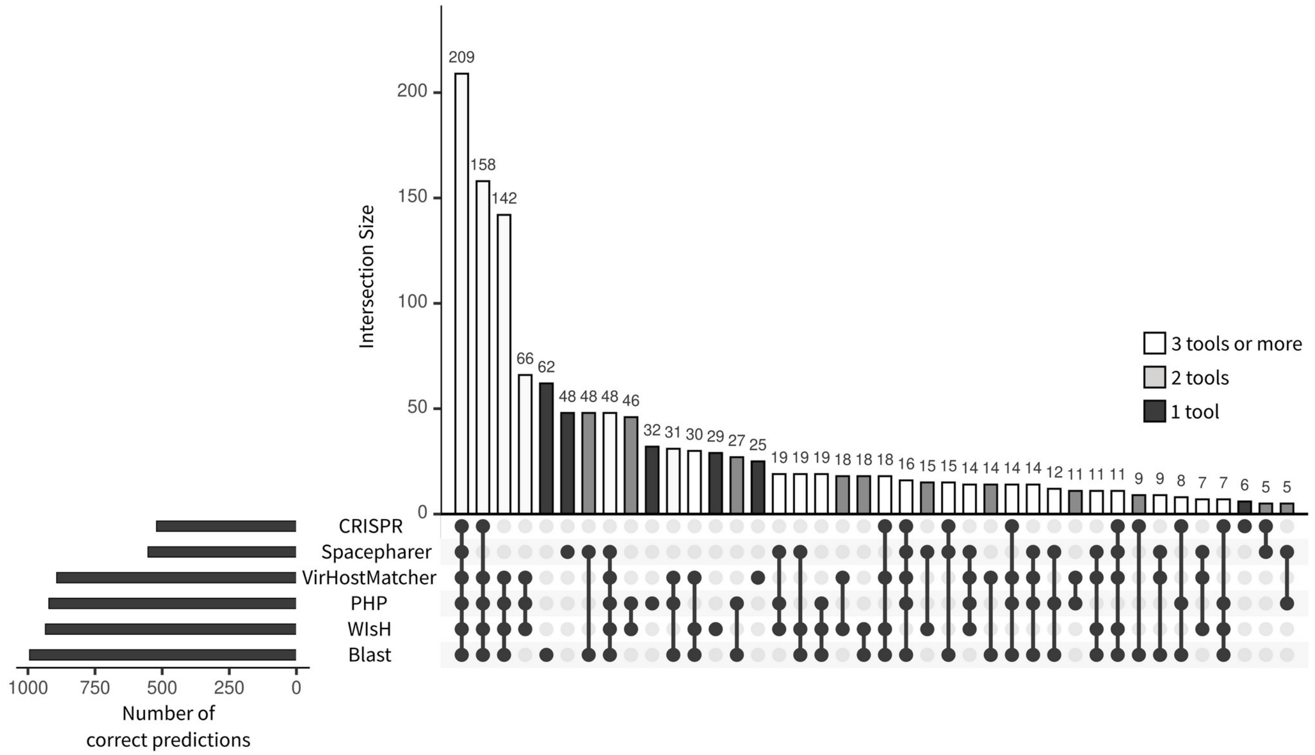


**Supplementary Figure S4. Overlap between host-based tools for individual viruses.** For each host-based tool included in the benchmark (see Fig. 1), the overlap in terms of input sequence for which a correct prediction was obtained is presented here as an upset plot. The intersection size represents the number of phages with correct prediction using the combination of methods indicated at the bottom. This number is also indicated above each bar, and the bar color indicates the number of tools included in the combination.

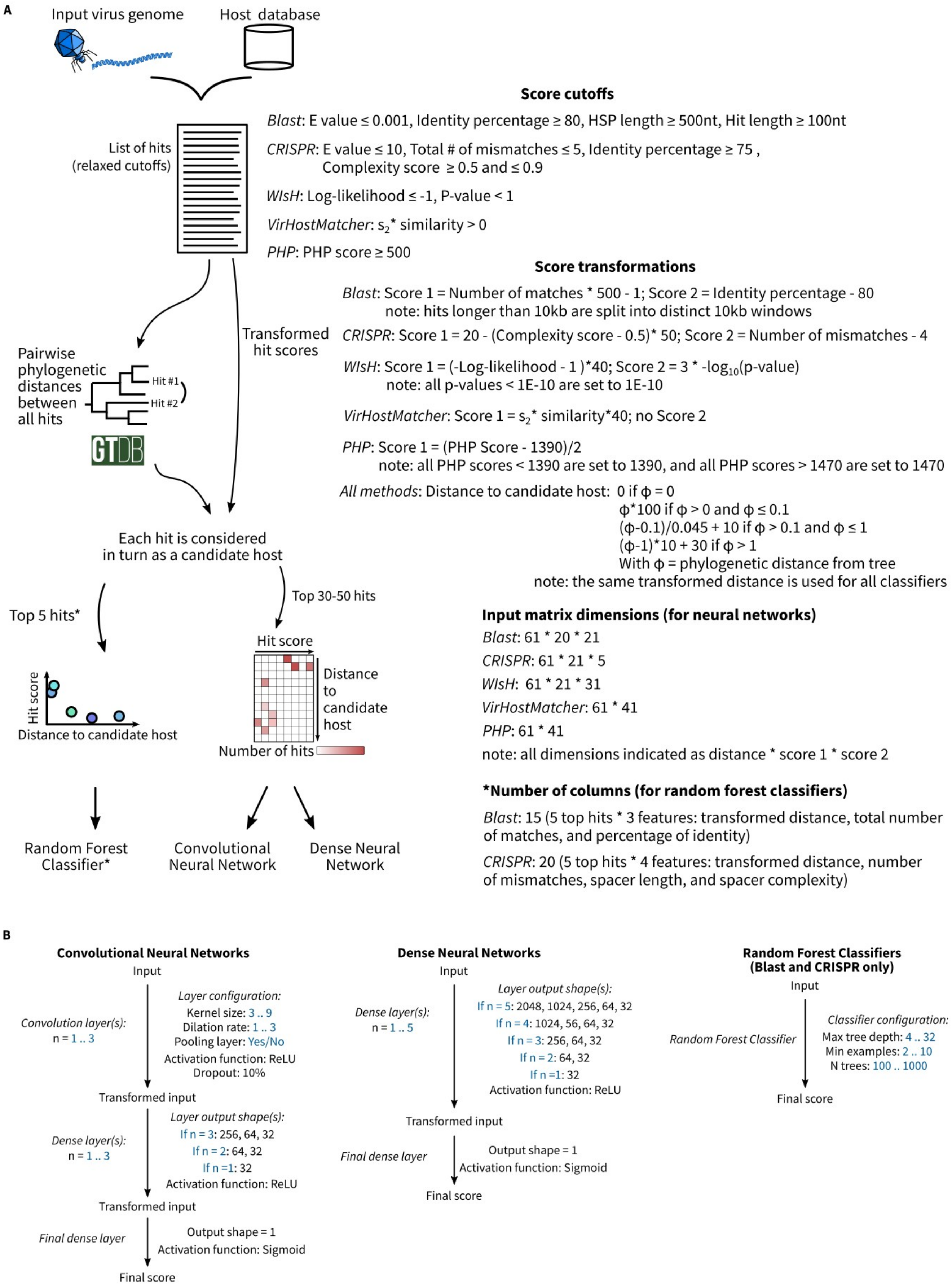
A
Input virus genome
Host database
List of hits
(relaxed cutoffs)
Score cutoffs
Blast: E value ≤ 0.001, Identity percentage ≥ 80, HSP length ≥ 500nt, Hit length ≥ 100nt
CRISPR: E value ≤ 10, Total # of mismatches ≤ 5, Identity percentage ≥ 75 , Complexity score ≥ 0.5 and ≤ 0.9
WIsH: Log-likelihood ≤ -1, P-value < 1
VirHostMatcher: $s_2$* similarity > 0
PHP: PHP score ≥ 500
Score transformations
Blast: Score 1 = Number of matches * 500 - 1; Score 2 = Identity percentage - 80
note: hits longer than 10kb are split into distinct 10kb windows
CRISPR: Score 1 = 20 - (Complexity score - 0.5)* 50; Score 2 = Number of mismatches - 4
WIsH: Score 1 = (-Log-likelihood - 1 )*40; Score 2 = 3 * -$\log_{10}$(p-value)
note: all p-values < 1E-10 are set to 1E-10
VirHostMatcher: Score 1 = $s_2$* similarity*40; no Score 2
PHP: Score 1 = (PHP Score - 1390)/2
note: all PHP scores < 1390 are set to 1390, and all PHP scores > 1470 are set to 1470
All methods: Distance to candidate host: 0 if ϕ = 0
ϕ*100 if ϕ > 0 and ϕ ≤ 0.1
(ϕ-0.1)/0.045 + 10 if ϕ > 0.1 and ϕ ≤ 1
(ϕ-1)*10 + 30 if ϕ > 1
With ϕ = phylogenetic distance from tree
note: the same transformed distance is used for all classifiers
Pairwise phylogenetic distances between all hits
Hit #1
Hit #2
GTDB
Transformed hit scores
Each hit is considered in turn as a candidate host
Top 5 hits*
Top 30-50 hits
Hit score
Distance to candidate host
Hit score
Distance to candidate host
Number of hits
Random Forest Classifier*
Convolutional Neural Network
Dense Neural Network
Input matrix dimensions (for neural networks)
Blast: 61 * 20 * 21
CRISPR: 61 * 21 * 5
WIsH: 61 * 21 * 31
VirHostMatcher: 61 * 41
PHP: 61 * 41
note: all dimensions indicated as distance * score 1 * score 2
*Number of columns (for random forest classifiers)
Blast: 15 (5 top hits * 3 features: transformed distance, total number of matches, and percentage of identity)
CRISPR: 20 (5 top hits * 4 features: transformed distance, number of mismatches, spacer length, and spacer complexity)
B
Convolutional Neural Networks
Input
Convolution layer(s):
n = 1 .. 3
Layer configuration:
Kernel size: 3 .. 9
Dilation rate: 1 .. 3
Pooling layer: Yes/No
Activation function: ReLU
Dropout: 10%
Transformed input
Dense layer(s):
n = 1 .. 3
Layer output shape(s):
If n = 3: 256, 64, 32
If n = 2: 64, 32
If n =1: 32
Activation function: ReLU
Transformed input
Final dense layer
Output shape = 1
Activation function: Sigmoid
Final score
Dense Neural Networks
Input
Dense layer(s):
n = 1 .. 5
Layer output shape(s):
If n = 5: 2048, 1024, 256, 64, 32
If n = 4: 1024, 56, 64, 32
If n = 3: 256, 64, 32
If n = 2: 64, 32
If n =1: 32
Activation function: ReLU
Transformed input
Final dense layer
Output shape = 1
Activation function: Sigmoid
Final score
Random Forest Classifiers
(Blast and CRISPR only)
Input
Random Forest Classifier
Classifier configuration:
Max tree depth: 4 .. 32
Min examples: 2 .. 10
N trees: 100 .. 1000
Final score

**Supplementary Figure S5. Schematic of the data transformation and classifier architectures used in iPHoP**. A. Summary of the cutoff and metrics used for each host-based tool considered in iPHoP (see Table S1). B. Overview of the three different types of classifiers evaluated in iPHoP. The different parameters optimized using the Optuna framework are highlighted in blue. For varying numbers of layers, the same parameters were optimized for each layer, but each was optimized separately, i.e., the parameters values were independent between the different layers.

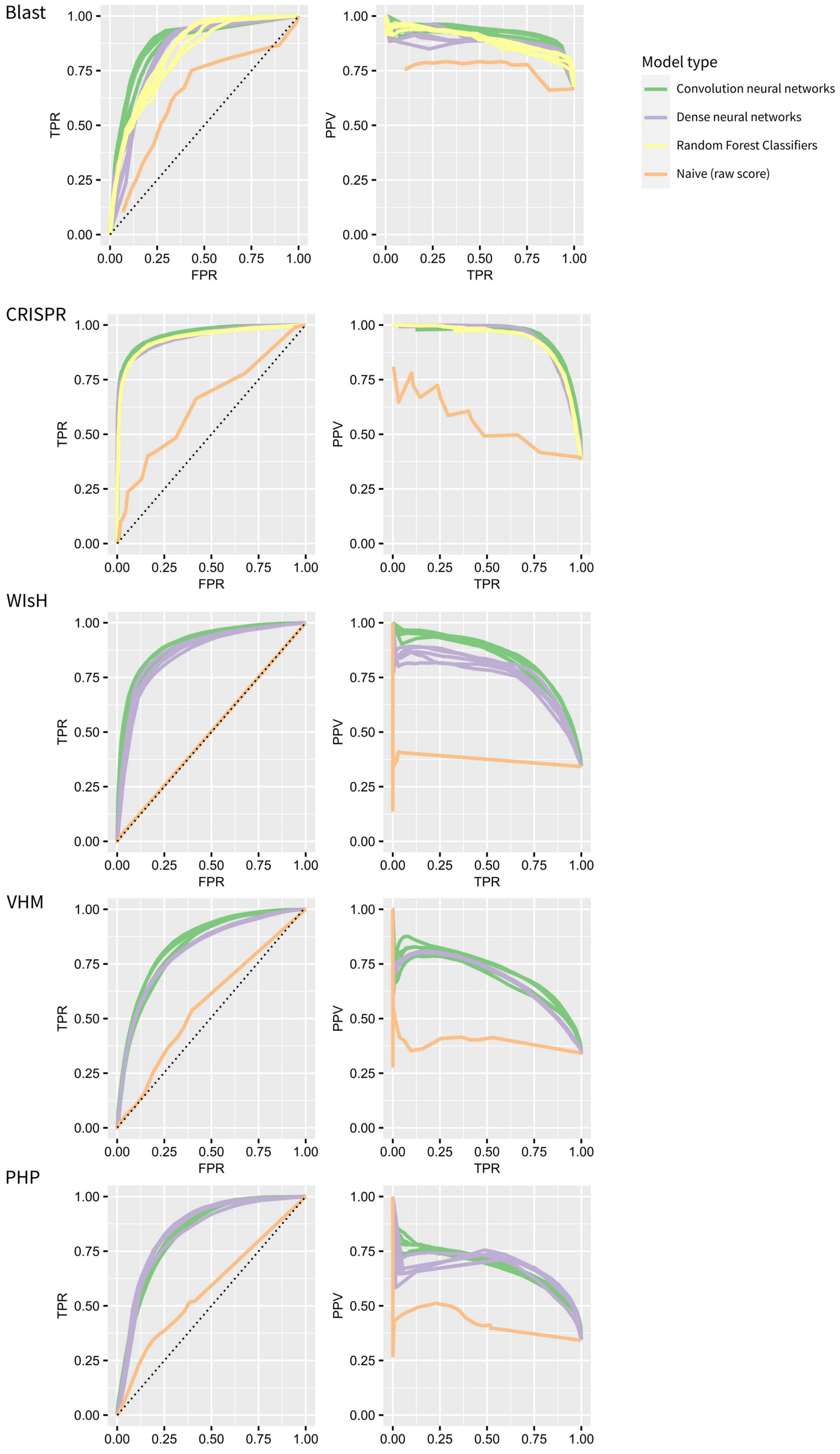

Blast
CRISPR
WIsH
VHM
PHP
TPR
FPR
PPV
0.00
0.25
0.50
0.75
1.00
Model type
Convolution neural networks
Dense neural networks
Random Forest Classifiers
Naive (raw score)

**Supplementary Figure S6. ROC and Precision-Recall curves for single-tool classifiers**. For each host-based tool the ROC curves (left) and Precision-Recall curves (right) based on the test dataset are presented for the 5 best classifiers of each type, and compared to the “naive” approach, i.e. best hit based on the raw score. TPR: True Positive Rate. FPR: False Positive Rate. PPV: Positive Predictive Value. The 1-to-1 line is indicated as a dashed black line on the ROC curves. Random Forest Classifiers were only evaluated for Blast and CRISPR approaches.

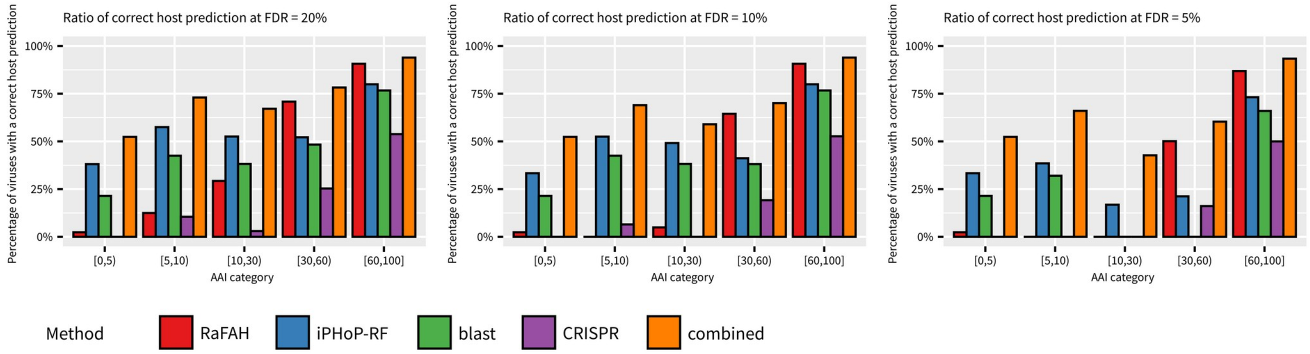


**Supplementary Figure S7. Percentage of correct host predictions obtained for viruses with different degrees of "novelty".** The number of correct host predictions was evaluated for 3 different score cutoffs corresponding to 20%, 10%, and 5% estimated FDR (False Discovery Rate). Input viruses were classified into 5 categories (x-axis) based on their AAI (Average Amino Acid Identity) to the closest reference phage genome. The number of correct host predictions is indicated for each iPHoP classifier (see Fig. 3A), and for the composite score considering all classifiers ("combined").

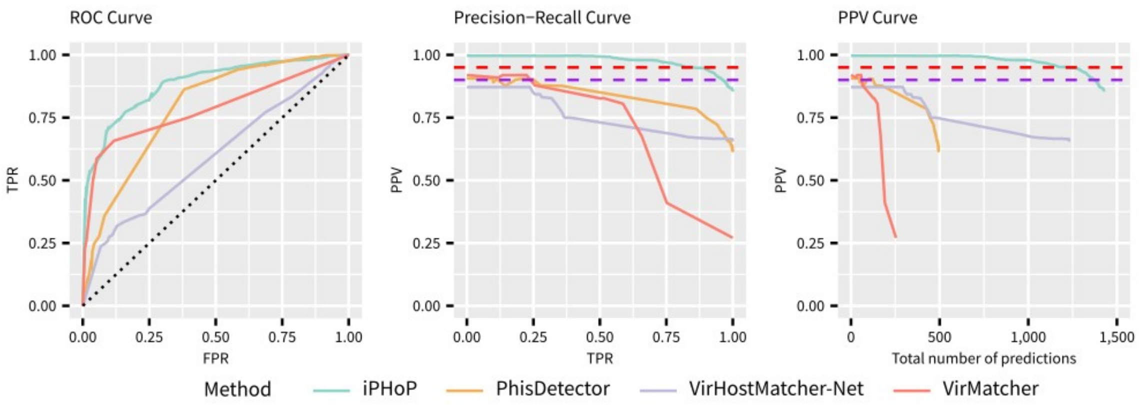


**Supplementary Figure S8. Comparison of different integrated host prediction tools, including iPHoP, on the test dataset.** Standard Receiver Operating Characteristic (left) and Precision Recall (middle) curves for the 4 integrated host prediction approaches compared. To take into account the number of predictions provided by each tool, a third plot (right panel) indicates the positive predictive value (y-axis) when considering an increasing number of predictions (x-axis). To obtain this, cutoffs were progressively lowered to include an increasing number of predictions for each tool, and prioritize the highest confidence ones, i.e. starting with the highest PPV possible. For the ROC curve, a 1-to-1 line is indicated with a dashed black line. For the Precision Recall and PPV curves (middle and right panels), the red and purple dashed lines indicate 5% and 10% False Discovery rates, respectively.

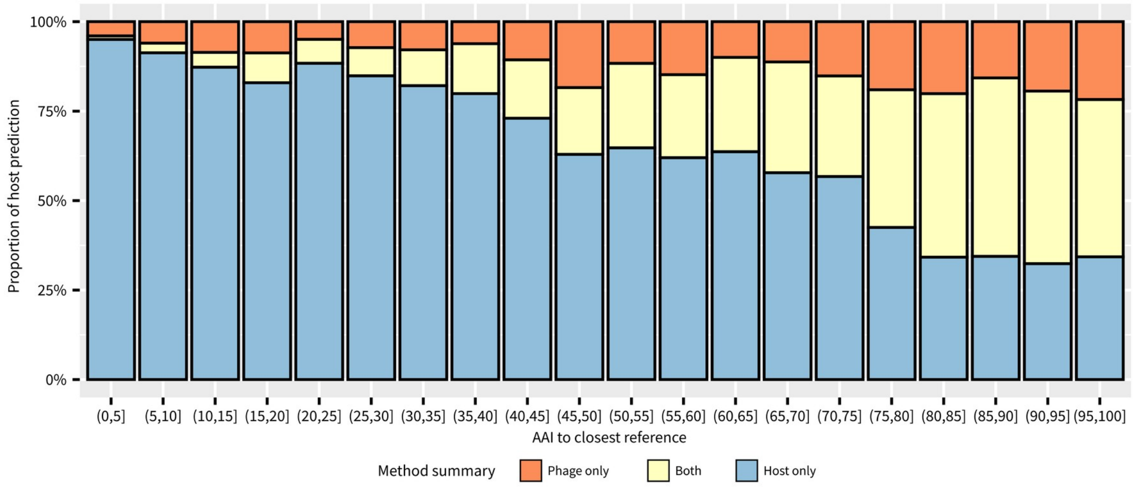


**Supplementary Figure S9. Type of host prediction obtained for high-quality IMG/VR v3 genomes with different degrees of "novelty".** High-quality genomes from the IMG/VR v3 database for which a host prediction was obtained with iPHoP (score $\geq$ 90) were binned based on the average amino acid identity (AAI) to the closest reference in NCBI RefSeq Virus r203 (x-axis). Predictions entirely based on host-based tools are indicated as "Host only", predictions exclusively based on RaFAH are indicated as "Phage only", and predictions where both types of tools were consistent and with score $\geq$ 90 are listed as "Both".

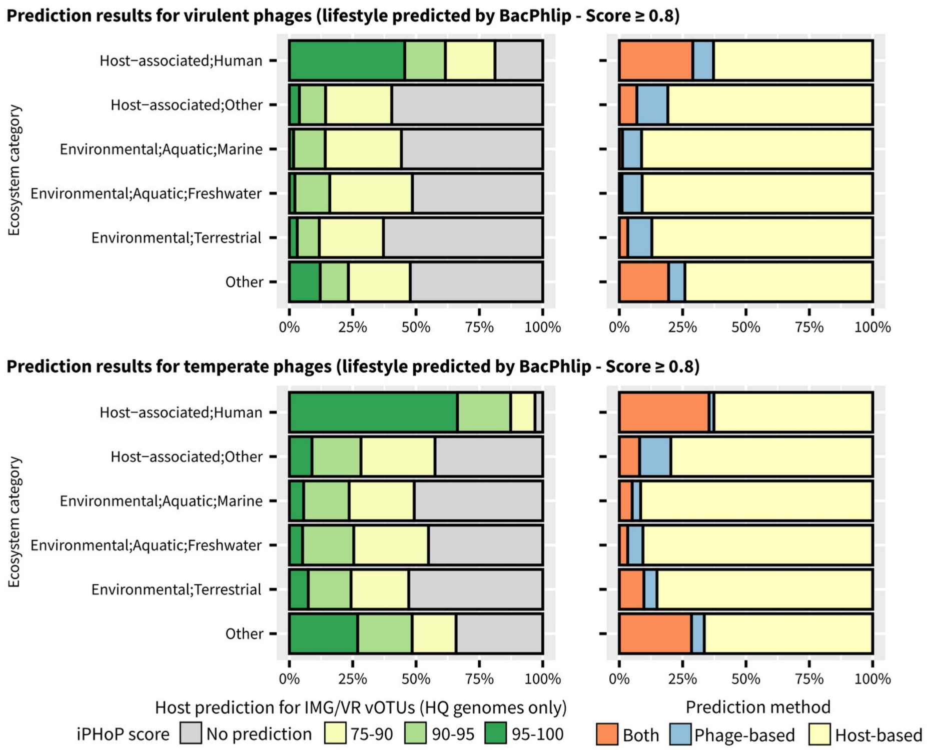


**Supplementary Figure S10. Breakdown of iPHoP host predictions for high-quality IMG/VR v3 genomes assigned as virulent (top) or temperate (bottom).** Similar as Fig. 4A and 4B, the left panel shows the distribution of the best score provided by iPHoP for the corresponding subset of IMG/VR v3 quality genome (top: virulent, bottom: temperate), organized by ecosystem. For each vOTU, the best score from iPHoP was considered if ≥ 75, or the vOTU was considered as not having a predicted host. The right panel shows the the type of signal used to achieve host prediction with a score ≥ 90. “Host-based” includes all 5 host-based tools, while “Phage-based” includes predictions obtained with RaFAH. “Both” includes consistent predictions obtained with RaFAH and at least one host-based tool. Temperate and virulent phages were identified via BACPHLIP[34] with a minimum score of 0.8 and based on genome annotation (see Methods).

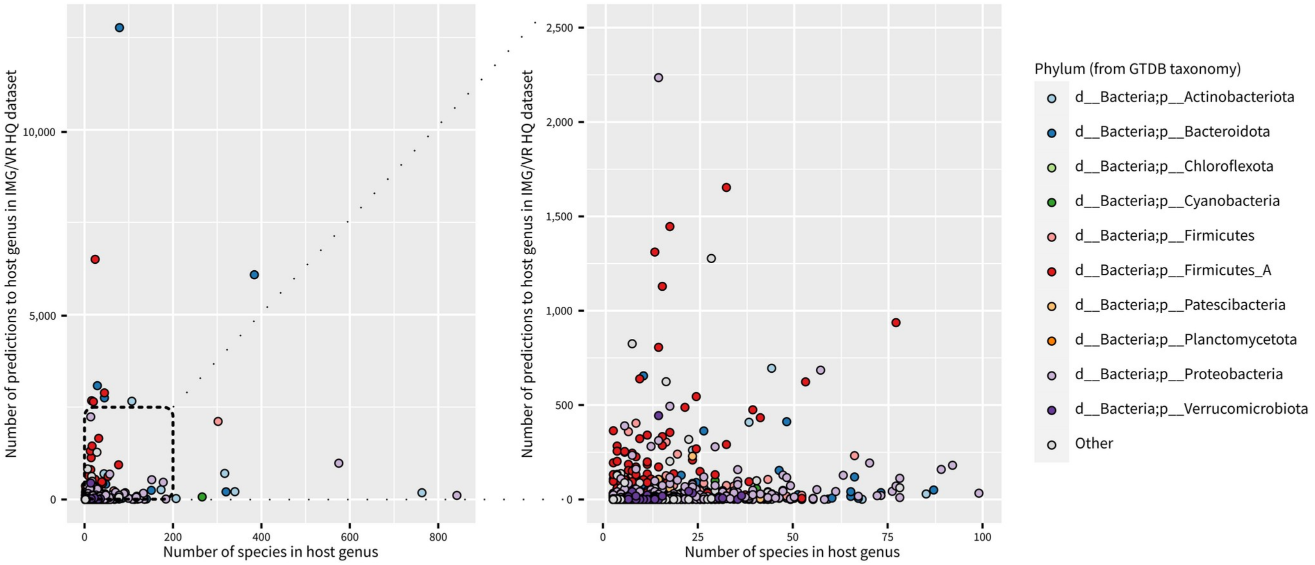


**Supplementary Figure S11. Number of species and iPHoP prediction per host genus.** Each dot represents a host genus with at least 2 species, with the x-axis reflecting the total number of species in the genus, and the y-axis reflecting the total number of IMG/VR v3 HQ sequences predicted to infect this host genus with a score ≥ 90. Host genera and species were obtained from the GTDB database[34]. The right panel presents a zoomed-in version of the area highlighted with dashed black lines in the left panel.
